# Enhancing metabarcoding efficiency and ecological insights through integrated taxonomy and DNA reference barcoding: a case study on beach meiofauna

**DOI:** 10.1101/2024.04.01.587521

**Authors:** Jan-Niklas Macher, Alejandro Martínez, Sude Çakir, Pierre-Etienne Cholley, Eleni Christoforou, Marco Curini Galletti, Lotte van Galen, Marta García-Cobo, Ulf Jondelius, Daphne de Jong, Francesca Leasi, Michael Lemke, Iñigo Rubio Lopez, Nuria Sánchez, Martin Vinther Sørensen, M. Antonio Todaro, Willem Renema, Diego Fontaneto

## Abstract

Molecular techniques like metabarcoding, while promising for exploring diversity of communities, are often impeded by the lack of reference DNA sequences available for taxonomic annotation. Our study explores the benefits of combining targeted DNA barcoding and morphological taxonomy to improve metabarcoding efficiency, using beach meiofauna as a case study. Beaches are globally important ecosystems and are inhabited by meiofauna, microscopic animals living in the interstitial space between the sand grains, which play a key role in coastal biodiversity and ecosystem dynamics. However, research on meiofauna faces challenges due to limited taxonomic expertise and sparse sampling. We generated 775 new cytochrome c oxidase I DNA barcodes from meiofauna specimens collected along the Netherlands’ west coast and combined them with the NCBI GenBank database. We analysed alpha and beta diversity in 561 metabarcoding samples from 24 North Sea beaches, a region extensively studied for meiofauna, using both the enriched reference database and the NCBI database without the additional reference barcodes. Our results show a 2.5-fold increase in sequence annotation and a doubling of species-level Operational Taxonomic Units (OTUs) identification when annotating the metabarcoding data with the enhanced database. Additionally, our analyses revealed a bell-shaped curve of OTU richness across the intertidal zone, aligning more closely with morphological analysis patterns, and more defined community dissimilarity patterns between supralittoral and intertidal sites. Our research highlights the importance of expanding molecular reference databases and combining morphological taxonomy with molecular techniques for biodiversity assessments, ultimately improving our understanding of coastal ecosystems.

## Introduction

Metabarcoding has become a standard technique for the study of biological communities and a valuable tool for understanding biodiversity and ecology of species and communities (Ficetola & Taberlet, 2023; Taberlet et al., 2012). Despite technological advances, the efficiency of metabarcoding studies and their usability for ecological analyses depends on the completeness of reference sequence databases. These databases, such as NCBI GenBank (Benson et al., 2012) and Barcode of Life (BOLD; (Ratnasingham & Hebert, 2007)) contain reference DNA barcodes (Hebert et al., 2003) generated from individual, taxonomically identified specimens that serve as reference against which the metabarcoding sequences can be compared and identified. The availability of such references greatly varies across taxonomic groups and ecosystems, remaining particularly poor for species-rich but understudied taxa and regions. (Keck et al., 2023; Leasi et al., 2018; Leray & Knowlton, 2015; A. M. Weigand & Macher, 2018). A promising approach to mitigate this issue lies in the targeted collection, identification, and sequencing of a high number of specimens, followed by their addition to molecular reference databases. This method, sometimes implemented through taxonomic expert workshops, has been successfully employed across diverse taxonomic groups and ecosystems (Behrens-Chapuis et al., 2021; Creedy et al., 2020; De Souza Amorim et al., 2023; Dugal et al., 2022; Emerson et al., 2023; Zhou et al., 2011).

Using beach meiofauna as a case study, we investigate the effect of targeted sampling and expert identification on improving metabarcoding efficiency by increasing the completeness of reference libraries. Beach meiofauna comprises small animals inhabiting the interstitial spaces between sand grains. They form highly diverse communities, providing important ecosystem services and are valuable for the understanding of biodiversity and ecosystem dynamics of coastal areas (Felix et al., 2016; Schratzberger & Ingels, 2018). Beaches are one of the Earth’s most dynamic environments, which is threatened by sea level rise and other human impacts (Lansu et al., 2024; Schlacher et al., 2007). Studying meiofauna is challenging due to their generally microscopic size and the scarcity of taxonomists for most of the world’s geographic regions and taxonomic groups (Leasi & Norenburg, 2014; Zeppilli et al., 2015). This taxonomic impediment means many beach meiofauna species remain undescribed even in intensively sampled areas (Curini-Galletti et al., 2012; Martinez et al., 2023). Advancements in DNA metabarcoding have drastically improved our ability to detect species in complex communities (Ficetola & Taberlet, 2023), and after pioneering studies demonstrating the use of DNA metabarcoding for marine meiofauna (Creer et al., 2010; Fonseca et al., 2010), the technique now allows for more detailed studies and gaining a better understanding of the impact of environmental factors and stressors on meiofauna (Atherton & Jondelius, 2020; Gielings et al., 2021; Martínez et al., 2020).

Here we introduce a dataset comprising 775 newly generated cytochrome c oxidase I reference sequences (“DNA barcodes”) derived from meiofauna specimens collected along the Netherlands’ west coast. This region, a part of the southern North Sea, is one of the best-studied areas in marine meiofaunal studies (Germán Rodríguez, 2004; Gray & Rieger, 1971; Kotwicki et al., 2005). By analysing a total of 576 metabarcoding samples, we study the influence of enhanced reference databases on inferred species richness and community composition patterns, both on a local (across the intertidal zone) and regional (spanning 650 km of the North Sea Coast) scales. By demonstrating major improvements in identification of meiofauna in metabarcoding data through reference database enhancement, our study highlights the critical need for combining taxonomic expertise with reference sequencing even in well-studied areas. Due to the massive improvement demonstrated by the use of a local reference library in one of the best-studied areas in the world, the improvement that can be obtained in poorly studied, species-rich areas will be much stronger. We seek to inspire similar initiatives in biodiversity research and advocate for combining traditional taxonomic methods and modern genetic techniques to improve the understanding of biodiversity in important, but often overlooked taxonomic groups and ecosystems.

## Materials and Methods

### Sampling for reference barcoding

Meiofauna specimens were collected during two taxonomic workshops held at Naturalis Biodiversity Center in Leiden, the Netherlands, in May/June 2022 and July/August 2023. Samples were collected from 20 locations along the Dutch West Coast, by either taking sediment cores to a depth of 10cm, by filtering coastal groundwater and sand through dug holes, or by scraping hard substrates, depending on the targeted taxonomic group. The samples were transported to the Naturalis laboratory for meiofauna extraction using decantation through a 40µm sieve after anesthetization with isosmotic MgCl_2_ (Somerfield & Warwick, 2013). For a detailed list of all sample locations and sample types, see supplementary table S1. After extraction, meiofauna specimens were identified to the lowest possible taxonomic rank using stereo and light microscopes. Specimens were transferred to PCR plates and submitted to the DNA extraction pipeline described below.

### DNA extraction and amplification for reference barcoding

The DNA extraction process for meiofauna specimens was performed using the Macherey-Nagel (Düren, Germany) NucleoSpin tissue kit on the KingFisher (Waltham, USA) robotic platform, following the manufacturer’s protocol. After extraction, PCRs were performed with Geller COI primers (Geller et al., 2013), targeting the 658 base-pair-long Folmer fragment of the mitochondrial cytochrome c oxidase I gene, which is the commonly used DNA barcode for animals (Folmer et al., 1994; Hebert et al., 2003). Samples were sequenced on the Oxford Nanopore GridION platform (Oxford Nanopore Technologies, Oxford, United Kingdom). The protocol was as follows: Each PCR reaction contained 10.2 µl of MiliQ water, 4 µl of 5X PCR buffer (Qiagen; Hilden, Germany), 0.8 µl of 10 mg/ml BSA (Promega, Madison, Wisconsin, United States), 1 µl of 10 picomolar/µl primers, 0.4 µl of 2.5mM dNTPs, 0.4 µl of 5U/µl Phire II Taq polymerase (Thermo Fisher, Waltham, Massachusetts, United States), and 1 µl template DNA. PCR started with an initial denaturation of 30 seconds at 98 °C, followed by 35 cycles of 5 seconds denaturation at 98 °C, 10 seconds annealing at 50 °C, 15 seconds elongation at 72 °C, and a final extension of 5 minutes at 72 °C. Each PCR included a negative control using Milli-Q water (Merck; Rahway, New Jersey, United States) instead of template DNA. We cleaned the PCR products using AmPure magnetic beads (Brea, California, United States) with a ratio of 0.9:1. Subsequently, a second PCR was performed to individually label the samples, with 2.5 µl of ONT barcode primers, 5 µl of LongAmp Taq 2x master mix (New England Biolabs, Ipswich, Massachusetts, United States), and 2.5 µl of the PCR product as template. The PCR protocol for the second amplification was as follows: an initial denaturation of 3 minutes at 95 °C, followed by 15 cycles of 15 seconds denaturation at 95 °C, 15 seconds of annealing at 65 °C, 50 seconds of elongation, and a final extension step of 3 minutes at 65 °C. The success was checked using the TapeStation platform. The samples were then pooled at equimolar concentrations to achieve a final concentration of approximately 200 femtomolar. This was followed by a purification step using a 0.7:1 bead cleanup, targeting amplicons of 700 bp length. Finally, the purified DNA pools were eluted in 11 µl of nuclease-free water and their concentration was quantified using the Tape station (D5000 kit).

### Sequencing and bioinformatics for reference barcoding

Sequencing was conducted using the Oxford Nanopore GridION sequencer on two FLO-MIN112 flow cells, with the SQK-NBD112.24 sequencing kit. The Basecalling was done with MinKNOW (v23.04.5), the run duration was set to 72H, and super accuracy basecalling was selected. The demultiplexing was performed with Guppy barcoder (v6.4.6). The consensus calling consisted of several steps combined together in a Snakemake (Mölder et al., 2021) pipeline: First, the reads (containing primers at both ends) were filtered by size (>=558, <= 758) and quality (>=10), and then reoriented with Cutadapt (v4.5, max error rate 20%, 80% coverage), which also removed flanking sequences. Then consensus sequences were generated using NGSpeciesID v0.3.0 (Sahlin et al., 2021) with Medaka polishing (v1.8.0, r104_e81_sup_g5015 model). A final round of primer sequence trimming was performed with Cutadapt. Following this, multi-fasta files containing consensus sequences were written by using a custom script. Quality control and visualisation of the processed FASTQ files was conducted using NanoPlot (De Coster et al., 2018) and MultiQC (Ewels et al., 2016). All resulting sequences underwent manual curation in Geneious Prime (version 2023.2) and were searched against existing references in NCBI GenBank using BLASTn (Ye et al., 2006). All scripts used for processing of Nanopore data are available on GitHub: https://gitlab.com/arise-biodiversity/sequencing/arise-barcoding-pipeline/-/tree/1a2fa544615be54dccb7136ca20d3664ba85d467.

### Sampling and environmental variable measurement for metabarcoding

We collected meiofauna samples for metabarcoding from 24 sandy beaches along the Dutch and German Coast during the summers of 2021 and 2022. See Figure 1 for a map, and Supplementary Table S2 for the coordinates of the sampled beaches. Sampling was conducted at maximum low tide. At each beach, we sampled along three parallel transects, with eight sampling sites each. The first sampling site was located at the foot of the dunes, the second midway between the dunes and the high-tide line, and the remaining six samples evenly spaced from the high-tide line to the low-tide line. Following McLachlan’s classification (McLachlan et al., 2018), most sampled beaches were tide-modified. Two beaches were wave dominated (Relative Tide Range Index (RTR) of <3), and two beaches were tide dominated (RTR >10). At each sampling site along the transects, two sediment cores were collected using sterile plastic syringes: one with a 5 cm diameter and 10 cm length (approximate volume of 200 ml), and another core with 1 cm diameter and 10 cm length (approximate volume of 8 ml). The smaller core was immediately transferred into a 50 ml Falcon tube, while the larger core was stored in a sterile 1-litre plastic bottle. On the beach, we extracted meiofauna from the larger core using the MgCl_2_ decantation method, by adding 500 ml of isosmotic MgCl_2_ solution to the sediment. After a 5-minute incubation, the sediment-MgCl_2_ mixture was swirled ten times, and the supernatant containing meiofauna was decanted through a 1 mm and 40 µm sieve cascade, a common practice in beach meiofauna studies (Castro et al., 2021; Haenel et al., 2017; Martínez et al., 2020). The meiofauna retained on the 40 µm sieve was then rinsed into sterile 15 ml Falcon tubes and preserved in 10 ml of 96% ethanol. All samples were subsequently transported to the Naturalis Biodiversity Centre laboratory and stored at -20°C until processing. The sediment from the smaller core was dried for grain size measurement using a LS13320 Particle Size Analyzer (Beckman-Coulter, USA).

**Figure 1.**
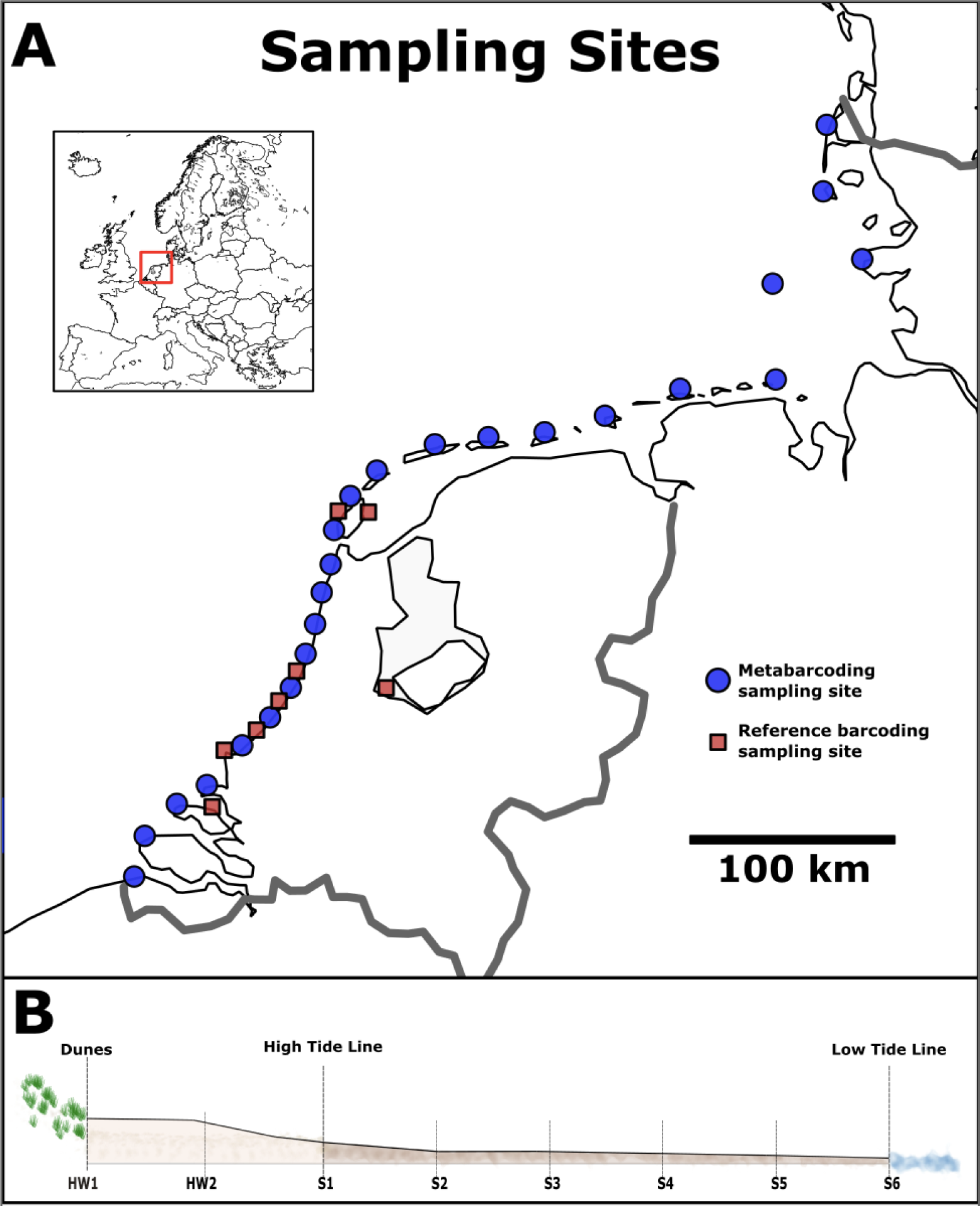
**A)** Map showing the location of the 24 beaches sampled for metabarcoding (blue circles) and locations sampled for reference barcoding (red squares). Note that reference barcoding sites in close proximity to each other are not shown separately due to the map scale. The mini map shows the locations of the sampling area in Europe. **B)** Schematic view of a beach showing the location of the eight sampling points per transect from dunes to the low tide line.

### DNA extraction, amplification and sequencing for metabarcoding

We extracted DNA from dried meiofauna samples after evaporating the ethanol at 50°C overnight in a sterile warming cabinet and transferring the dried samples to 2 ml Eppendorf tubes. DNA extraction was performed using the Macherey Nagel NucleoSpin Soil kit (Macherey Nagel, Düren, Germany) following the standard protocol including bead beating, but with an additional overnight Proteinase K digestion step (50 µl 250 µg/ml ProtK, Thermo Fisher Scientific, Waltham, USA) added to the lysis buffer provided with the kit to improve cell lysis, as done in previous studies on meiofauna (Martínez et al., 2020; A. M. Weigand & Macher, 2018).

For community metabarcoding, we amplified meiofauna DNA using a two-step PCR protocol with the widely-used LerayXT primers targeting a 313 base pair region of the mitochondrial cytochrome c oxidase I (COI) gene of a broad range of Eukaryota (Leray et al., 2013; Wangensteen et al., 2018). The first PCR reaction contained 11.7 µl mQ water, 2 µl Qiagen CL buffer (10x; Qiagen, Hilden, Germany), 0.4 µl MgCl2 (25 mM; Qiagen), 0.8 µl Bovine Serum Albumine (BSA, 10 mg/ml), 0.4 µl dNTPs (2.5 mM), 0.2 µl Qiagen Taq (5U/µl), 1 µl of each nextera-tailed primer (10 pMol/µl), and 2.5 µl of DNA template. PCR amplification involved an initial denaturation at 96°C for 3 minutes, followed by 30 cycles of denaturation for 15 seconds at 96°C, annealing at 50°C for 30 seconds, and extension for 40 seconds at 72°C, concluding with a final extension at 72°C for 5 minutes. We processed six negative controls (Milli-Q water, Merck, Kenilworth, USA) alongside the samples to check for potential contamination. After the first PCR, samples were cleaned with AMPure beads (Beckman Coulter, Brea, United States) at a 0.9:1 ratio according to the protocol to remove short fragments and primer dimers. The second PCR involved amplification with individually tagged primers, following the same protocol as above and using the PCR product from the first PCR as the template, but reducing the PCR cycle number to 10. We measured DNA concentrations using the Fragment Analyzer (Agilent Technologies, Santa Clara, CA, USA) with the High Sensitivity Kit and pooled samples equimolarly. The final library was cleaned with AMPure beads as described above and sent for sequencing on three Illumina MiSeq runs (2 × 300 bp read length) at Baseclear (Leiden, The Netherlands).

### Bioinformatic processing of metabarcoding data

The raw metabarcoding reads were processed using APSCALE (Buchner et al., 2022) with the following settings: maximum differences in percentage: 20; minimum overlap: 50, minimum read sequence length: 310 bp; maximum read length: 316 bp, minimum size to pool: 20 sequences. Sequences were clustered into both ESVs (setting: alpha 1, minimum 20 sequences) and OTUs, the latter with a sequence similarity threshold of 97%. To account for potential low-level contamination or tag jumping common on Illumina platforms (Schnell et al., 2015), we removed OTUs with an abundance of <0.03% of reads per sample, and OTUs that were present in less than 6 out of 561 samples (<1% occurrence). The taxonomic assignment was first performed using NCBI GenBank, and then with GenBank data expanded with the meiofauna COI barcodes generated as part of the taxonomic workshops in Leiden. Taxonomic ranks were assigned to OTUs using established identity thresholds: >97%: species, >95%: genus, >90%: family, >85% order (T.-H. Macher et al., 2023). OTUs that were assigned with less than 85% identity to a reference or identified as non-meiofauna taxa were excluded from further analyses. The three replicates per tidal level per beach were merged into one composite sample to account for potential variability within tidal levels, resulting in 190 composite samples.

### Analysing the increase in annotation efficiency by enhancing the reference database

We evaluated the efficiency of taxonomic assignment by calculating the percentage of reads and the number of Operational Taxonomic Units (OTUs) assigned to species, genus, family, order, and class when using NCBI GenBank for identification, and when expanding the database with the newly generated DNA barcodes. For this, we downloaded NCBI GenBank, conducted a nucleotide blast (blastn) for annotation, and then a second blastn annotation with NCBI GenBank enhanced with the newly generated sequences. All analyses were performed on the Naturalis Biodiversity Center High Performance Computing Cluster.

### OTU richness and community similarity across the intertidal zone

We calculated the OTU richness for each tidal level, both for the dataset annotated only with the NCBI database, and for the dataset annotated with both NCBI and the newly generated reference barcodes. We created point plots showing the increase in OTU number per taxonomic group and tidal level using the ggplot2 package in R, and used paired t-tests to test for significance in difference of OTU numbers. Further, we visualised the total number of OTUs at each tidal level using Cumming estimation plots using the ‘dabestr’ package, and used estimation statistics (Ho et al., 2019) to assess the mean difference between OTU numbers between the two datasets. We calculated the proportion of OTU numbers per taxonomic group and tidal level and computed stacked bar plots using ggplot2 in R. To analyse differences in taxonomic composition between the datasets at each tidal level, we applied chi-square tests, incorporating a Monte Carlo simulation for p-value estimation (simulate.p.value = TRUE) to address the small sample sizes and low expected frequencies.

We analysed the distribution of OTUs across the beach transect from dunes to the low tide line, both for the dataset annotated only with NCBI GenBank and the dataset annotated with NCBI and the new reference barcodes. Tidal levels were ordered categorically from dune level (HW1) to low water level (S6) to reflect the natural gradient. To quantify the relationship between OTU richness and both tidal levels and the Relative Tide Range Index (RTR), we employed generalised linear mixed-effects models (GLMMs). This approach enabled us to assess the statistical significance of linear and potentially nonlinear relationships affecting OTU richness across tidal levels and RTR, while also accounting for potential within-beach correlation in OTU richness. We tested for overdispersion and to ensure the model represented the underlying data structure and OTU count variability, we selected Poisson distribution for the dataset annotated with NCBI only, and a negative binomial distribution for the dataset annotated with NCBI and the new reference barcodes.

We calculated beta diversity using the Jaccard index, based on presence-absence data, and used the adonis2 PERMANOVA implemented in the ‘vegan’ package in R to assess the influence of the tidal level, i.e., the distance from the low tide, and the Relative Tide Range Index (RTR), i.e., the beach type, on community composition, using ‘Beach’ as a stratifying factor. Further, we calculated Non-metric Multidimensional Scaling plots (NMDS) for visualisation. Subsequently, we used a Mantel test to assess the correlation and significance of the similarity between these datasets.

### Impact of proximity to reference barcoding sites on meiofauna OTU richness in metabarcoding data

We tested whether the distance between sites sampled for reference barcoding, during the taxonomic workshops, and the sites sampled for metabarcoding significantly influenced the increase in number of OTUs identified as meiofauna in our metabarcoding dataset. To do this, we employed a linear modelling approach in R. We first calculated the geographic distances of metabarcoding sampling sites to the nearest site sampled for reference barcoding. Then, we constructed linear models to explore the relationship between the percentage increase in OTU richness and the distance to the nearest reference site. This was repeated for all eight tidal levels.

## Results

We structure the results into two main sections. First, we report the striking difference in taxonomic classification when integrating the newly generated COI barcodes into the reference library. Second, we examine the impact of this enriched reference library on ecological interpretations when analysing the same metabarcoding dataset.

### Reference DNA barcoding

We sequenced 775 mitochondrial cytochrome c oxidase I (COI) DNA barcodes of meiofaunal specimens following two taxonomic workshops in Leiden in the years 2022 and 2023. All specimens were identified at least on class level, with the exception of Platyhelminthes, which were classified at least to the subphylum level. For 643 specimens (82.97%), we also identified at least the order level, 490 specimens (63.23%) were identified at least on family level, 383 specimens (49.42%) on the genus level, and 164 (21.16%) were identified on the species level. The majority of barcoded specimens were Copepoda (143), followed by Chromadorea and Enoplea (Nematoda), with 137 and 116 specimens respectively. Rhabditophora (Platyhelminthes) and Polychaeta were present with 80 and 75 specimens, respectively, followed by Gastrotricha (63 specimens), Acoela (41), Clitellata (30), Rotifera (28), Arachnida (20), Collembola (14), Eutardigrada (7), Heterotardigrada (7), Palaeonemertea (4), Ostracoda (3), Branchiopoda (1), and Gnathostomulida (1). In addition, we found and sequenced four specimens of Malacostraca, and one Asteroidea larva. See supplementary table S3 for the complete taxonomic list of barcoded specimens and COI barcode sequences. The specimen list is also available as a GBIF dataset: https://doi.org/10.15468/gemfv4.

### Increase in annotation efficiency of metabarcoding data by enhancing the reference database

From the 561 samples analysed through metabarcoding, we retained 14,822,456 sequences and 566 OTUs after bioinformatic processing and quality filtering. Merging the three biological replicates per tidal level resulted in 190 composite samples. Using only NCBI GenBank for annotation, 4,633,286 sequences in 114 OTUs were assigned to meiofaunal taxa at least at the phylum level. Using NCBI GenBank in combination with the newly generated reference barcodes, 11,361,563 of sequences in 188 OTUs were identified as meiofauna. This corresponds to an increase of 145% in reads and an increase of 65% in OTUs annotated to meiofauna at least at the phylum level. The majority of non-meiofaunal reads were assigned to Bacillariophyta (2,652,322 reads, corresponding to 76,64% of non-meiofaunal reads). The number of OTUs assigned to order level increased by 63%, from 115 to 188. On the family level, we found an increase of 160%, from 53 to 138 OTUs. The number of OTUs assigned to at least genus level increased by 202%, from 40 to 121, and the number of OTUs annotated to species level increased by 205%, from 36 to 110.

The OTU number increased for all taxonomic groups except Branchiopoda (Arthropoda) and Pilidiophora (Nemertea), with the strongest increase observed in Copepoda (Arthropoda), Chromadorea and Enoplea (both: Nematoda). Commonly, we found the strongest increase in number of OTUs in the middle to lower intertidal zone (sampling sites S3 to S6), with the exception of Clitellata (Annelida), Arachnida, and Collembola (both: Arthropoda), for which we found the strongest increase in OTU numbers in the supralittoral zone (sampling sites HW1, HW2) and the upper intertidal zone (sampling sites S1, S2). Heterotardigrada OTUs were only found in the dataset annotated with NCBI combined with the new reference barcodes (Figure 2). See supplementary table S4 for paired t-test results.

**Figure 2:**
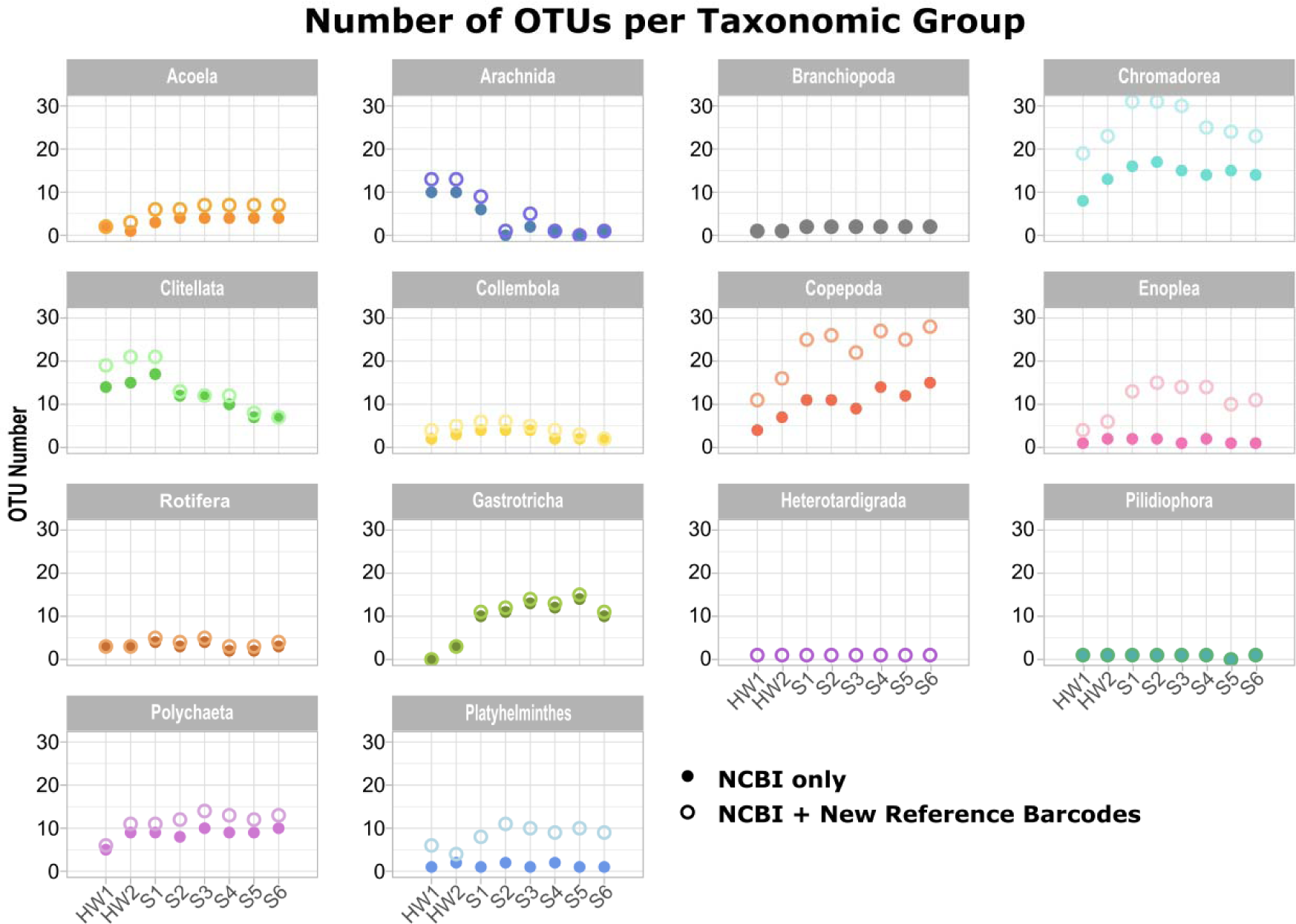
Increase of OTU number by taxonomic group and tidal level. Supralittoral sampling sites are labelled HW1 and HW2, and intertidal sampling sites are labelled S1 to S6. Filled circles indicate the number of OTUs identified using only NCBI GenBank for annotation of metabarcoding data, and open circles indicate the number of OTUs identified when combining NCBI GenBank with the new reference barcodes.

### Taxonomic composition in metabarcoding data

Comparing the dataset annotated with only NCBI reference barcodes and the database annotated with the new references, we found a consistent distribution of OTU proportions per taxonomic group across the intertidal zone. Enoplea (Nematoda) exhibited an increased proportion across all tidal levels when annotated with the expanded reference dataset, and a similar trend was observed for Platyhelminthes. Additionally, Heterotardigrada, while present in low proportions, were exclusively identified in the dataset enhanced with new reference barcodes (Figure 3). However, chi-square tests showed that overall proportions of taxonomic groups per tidal level did not differ significantly between the datasets (see supplementary table S5 for results).

**Figure 3:**
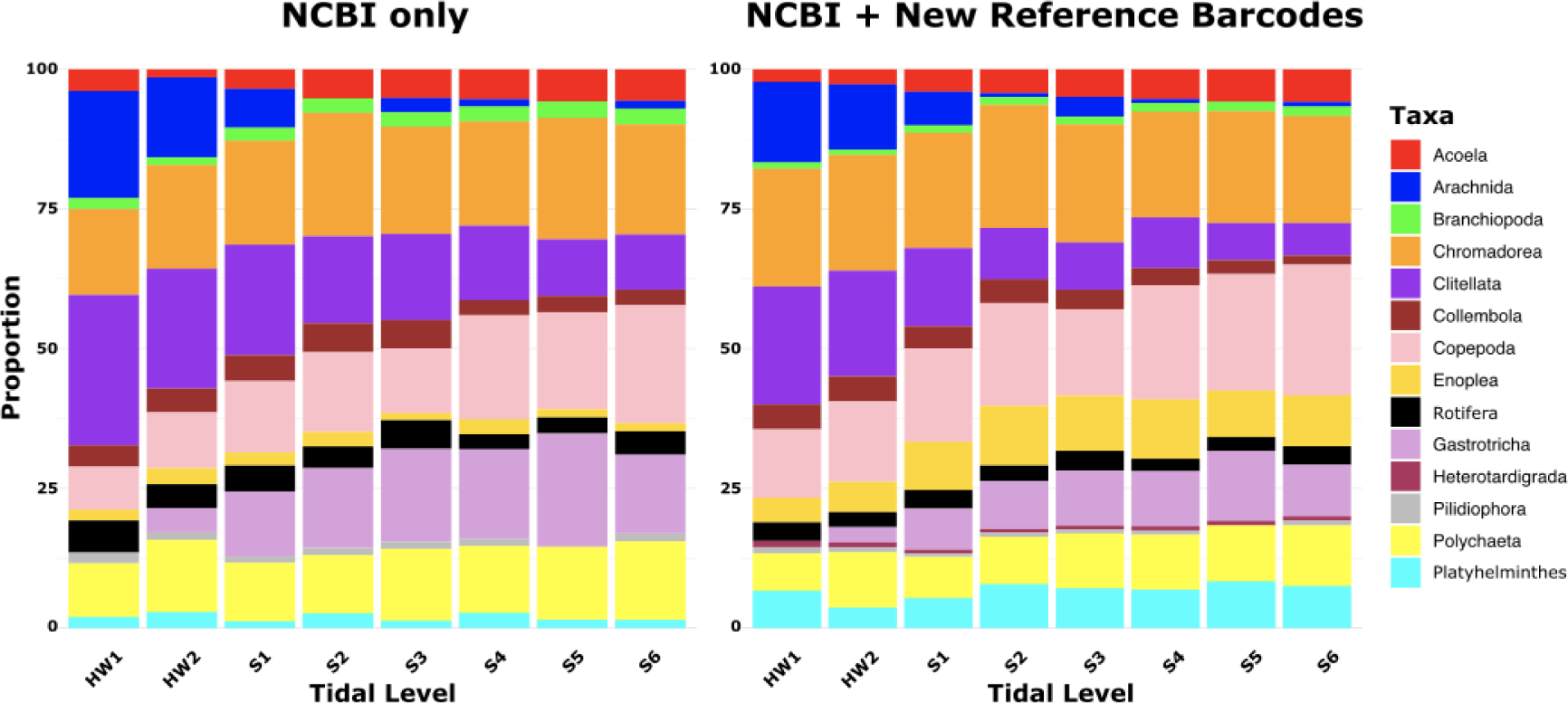
Stacked bar plots showing the proportions of meiofauna OTUs per taxonomic group across the beach transect. The left plot represents the dataset annotated using only NCBI reference barcodes and the right plot including both NCBI and newly generated reference barcodes. Each bar represents a tidal level (HW: sublittoral sites, S: intertidal sites), with the vertical stacking indicating the proportion of OTUs.

### OTU richness along beach transects

The enhancement of the molecular reference database significantly improved the assignment of Operational Taxonomic Units (OTUs) to meiofauna taxa for all tidal levels. The most notable increase was observed in tidal levels S3, S4, and S5, while the lowest increase was found for tidal levels HW1 and HW2. See Figure 4 for graphical representation and table 1 for statistical results.

**Figure 4:**
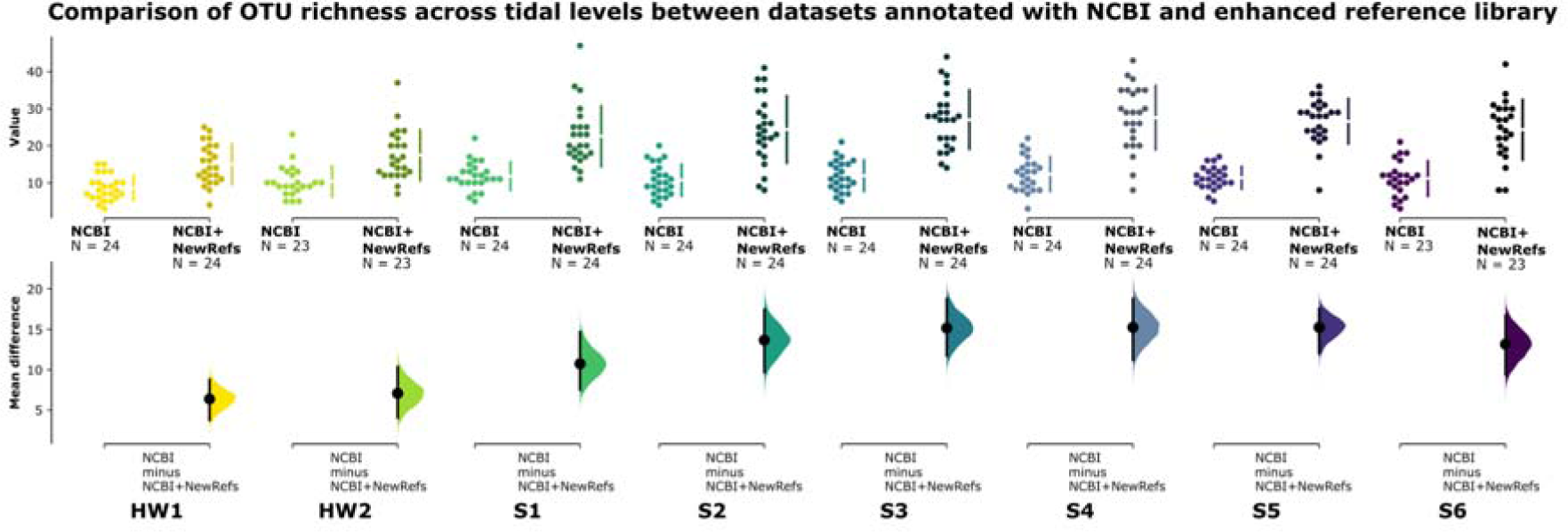
Cumming estimation plot showing the mean differences for OTU richness in the eight analysed tidal levels, between the dataset annotated with only NCBI reference data, and annotated with NCBI plus newly generated reference barcodes. The raw data is plotted on the upper axes; each mean difference is plotted on the lower axes as a bootstrap sampling distribution. Mean differences are depicted as dots; 95% confidence intervals are indicated by the ends of the vertical error bars.

**Table 1.** Unpaired Mean Differences, 95% Confidence Intervals (CI), and P Values from two-sided permutation t-tests comparing OTU richness per tidal levels between the dataset annotated with NCBI data only, and the dataset annotated with NCBI + newly generated reference barcodes. Results are based on 5000 bootstrap samples to assess the statistical significance of observed differences, with P values reflecting the probability of observing the effect sizes under the null hypothesis of no difference between groups.

| Tidal level | Unpaired Mean Difference | 95% CI | p value |
| --- | --- | --- | --- |
| HW1 | 6.38 | 3.75, 8.79 | <0.001 |
| HW2 | 7.09 | 4.04, 10.3 | <0.001 |
| S1 | 10.8 | 7.54, 14.7 | <0.001 |
| S2 | 13.7 | 9.71, 17.5 | <0.001 |
| S3 | 15.2 | 11.7, 18.8 | <0.001 |
| S4 | 15.2 | 11.2, 18.8 | <0.001 |
| S5 | 15.2 | 12.0, 17.5 | <0.001 |
| S6 | 13.2 | 9.39, 16.7 | <0.001 |

The analysis of OTU richness across beach transects, from dunes to the low tide line, revealed distinct patterns when comparing the datasets annotated only with NCBI GenBank and those enriched with new reference barcodes.

For the dataset annotated only with NCBI GenBank references, the GLMM revealed significant linear (p = 0.002) and quadratic (p = 0.003) effects of tidal levels on OTU numbers, indicating a non-linear distribution of OTU richness across the tidal gradient. The positive linear term suggests an increase in OTU numbers towards the middle and lower intertidal level, and the negative quadratic term indicates that this increase peaks at intermediate levels, rather than continuing linearly across the entire gradient. The random effects component attributed to beach location accounted for a variance of 0.037 (standard deviation: 0.1928), and the model’s AIC and BIC were 1044.2 and 1076.7, respectively. The analysis of the dataset annotated with NCBI and new reference barcodes showed a more pronounced non-linear (bell-shaped) pattern in OTU richness across tidal levels, with highly significant linear (p < 0.001) and quadratic (p < 0.001) relationships. This model accounted for a greater proportion of variability in OTU numbers, with a variance of 0.045 (standard deviation: 0.212) attributed to beach location, and the model’s AIC and BIC were 1278.8 and 1314.5, respectively. The influence of the Relative Tide Range Index (RTR) on OTU richness was not statistically significant in either dataset (p > 0.05), indicating that within the context of these models, RTR did not play a significant role in shaping OTU richness.

### Community composition

The analysis of community composition using Jaccard dissimilarity showed that in the dataset annotated only with NCBI data, the tidal level significantly influenced community composition (R² = 0.127, p < 0.001). The RTR, characterising beach type, also showed a significant but smaller effect (R² = 0.019, p = 0.001). The variability attributed to different Beaches was significant (R² = 0.273, p < 0.001) and had the highest explanatory value. A large proportion of variability in community composition remained unexplained (Residual R² = 0.581).

For the dataset annotated with both NCBI and new reference barcodes, we found a more substantial influence of the tidal level on community composition (R² = 0.155, p < 0.001). The effect of RTR remained similar to that observed in the NCBI-only dataset (R² = 0.018, p <0.001). The variability attributed to different Beaches was slightly more pronounced in this dataset (R² = 0.28, p < 0.001). The residual variability in this dataset was slightly lower (Residual R² = 0.547), indicating that the inclusion of new reference barcodes slightly improved the model’s explanatory power. See table 2.

**Table 2:** Summary of adonis2 PERMANOVA Results Comparing the Effects of Tidal Level, Relative Tide Range (RTR), and Beach on Inferred Community Composition Using Only NCBI and NCBI + New Reference Barcodes for Annotation of Metabarcoding Data.

| Test | Df | SumOfSqs | R<sup>2</sup> | F | Pr(>F) |
| --- | --- | --- | --- | --- | --- |
| **NCBI only** |  |  |  |  |  |
| Tidal level | 7 | 9.715 | 0.127 | 4.9756 | < 0.001 |
| RTR | 1 | 1.420 | 0.019 | 5.0896 | < 0.001 |
| Beach | 22 | 20.867 | 0.273 | 3.4004 | < 0.001 |
| Residual | 159 | 44.350 | 0.581 |  |  |
| **NCBI + new references** |  |  |  |  |  |
| Tidal level | 7 | 11.222 | 0.155 | 6.4542 | < 0.001 |
| RTR | 1 | 1.283 | 0.018 | 5.1640 | < 0.001 |
| Beach | 22 | 20.187 | 0.28 | 3.6943 | < 0.001 |
| Residual | 159 | 39.493 | 0.547 |  |  |

The Non-metric Multidimensional Scaling (NMDS) plots further underline these findings. In both datasets, samples collected from the highest tidal levels (HW1 and HW2) loosely cluster together, distinguishing them from samples taken from the middle to lower intertidal zone (S3 to S6). Samples from the upper intertidal or swash zone (S1 and S2) cluster between and within these two groups. The NMDS plot based on the dataset annotated with both NCBI and new references shows a clearer separation between the supralittoral sampling sites and the intertidal communities (see Figure 5).

**Figure 5:**
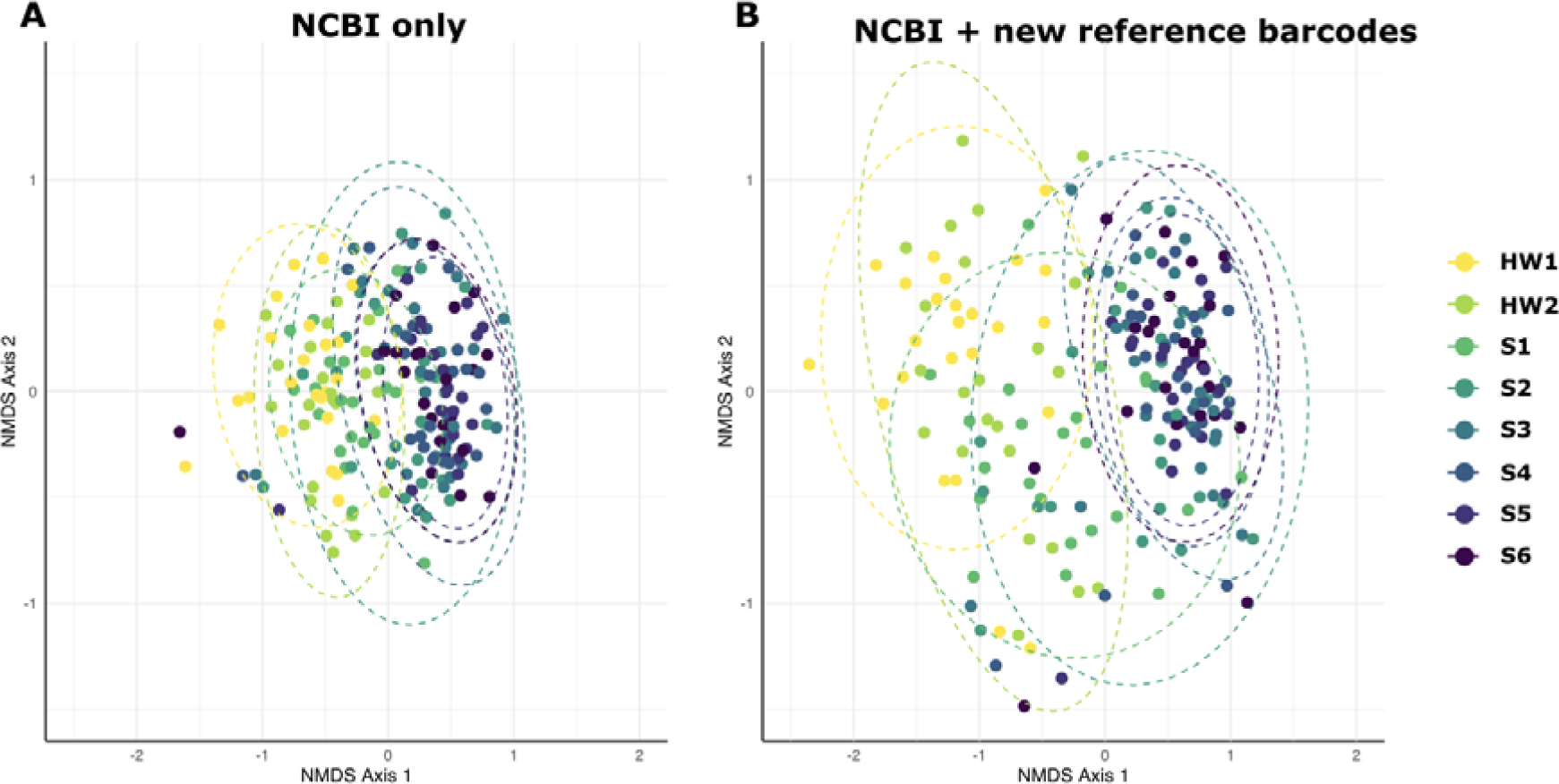
Non-metric Multidimensional Scaling (NMDS) ordination plots comparing community compositions across different tidal levels, based on Jaccard dissimilarity. A) Dataset annotated with NCBI reference data; B) Dataset annotated with NCBI and new reference barcodes. Points represent individual samples, coloured by their respective tidal level. Groups are outlined by 95% confidence ellipses.

The Mantel test results showed a strong and statistically significant correlation between the two distance matrices, with a Mantel statistic (r) of 0.7848 and a p-value of 0.001.

### Influence of distance from reference barcoding sites on the increase of identified meiofauna OTUs in metabarcoding data

We found negative correlations between the increase in identified meiofauna OTUs and the distance of the sampling site to a site sampled for reference barcoding. This was true for sampling sites in the tidal levels S6, S4, S3, S2, and S1. We observed a negative, but non-significant, trend for tidal level S5. We did not find correlations in the higher tidal levels HW1 and HW2 (Figure 6 and Table 3).

**Figure 6:**
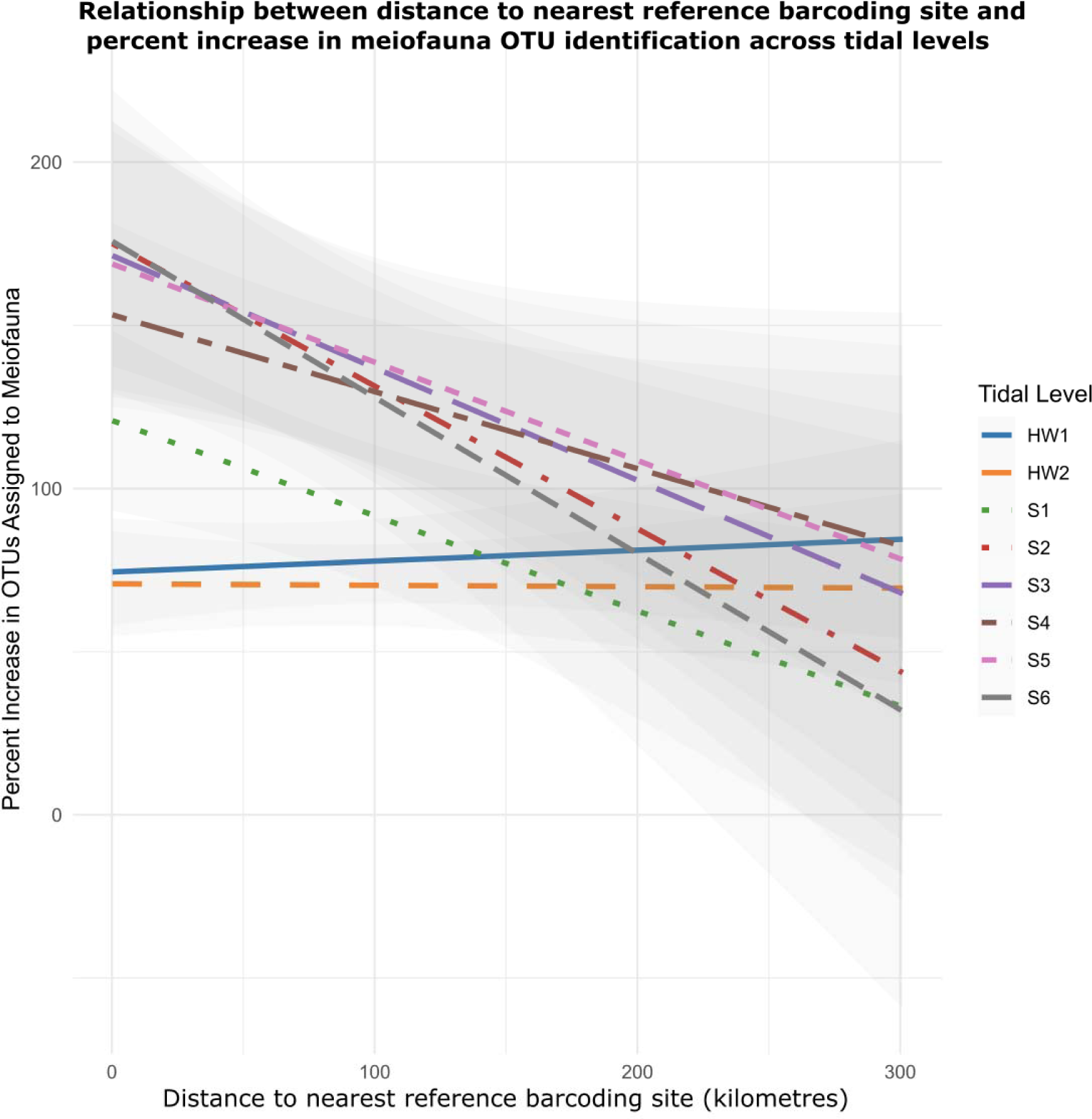
Linear relationship between the distance to the nearest reference site with newly generated reference barcodes (in kilometres) and the percent increase in Operational Taxonomic Units (OTUs) assigned to meiofauna. Linear regression lines in different colours and shaded confidence intervals show the trend for each tidal level.

**Table 3:** Linear Regression coefficients showing the change in increase in identified OTUs per kilometre distance to the nearest reference barcoding site for the eight analysed tidal levels. The coefficients indicate the magnitude and direction of the relationship, with negative values showing that the increase of OTU identifications slows down as distance to a reference barcoding sampling site increases.

| Tidal Level | Coefficient | p value |
| --- | --- | --- |
| HW1 | 0.00003379 | 0.595 |
| HW2 | -0.000001390 | 0.982 |
| S1 | -0.0002946 | 0.011 |
| **S2** | **-0.0004383** | **0.006** |
| **S3** | **-0.0003350** | **0.046** |
| **S4** | **-0.000232** | **0.044** |
| **S5** | -0.0003074 | 0.061 |
| **S6** | **-0.0004842** | **0.016** |

## Discussion

We tested the impact of enriching molecular reference databases on the efficiency of metabarcoding analyses, using beach meiofauna, an important yet frequently overlooked component of coastal biodiversity, as a case study. Our findings reveal an impressive improvement in meiofauna identification across all taxonomic levels following two targeted reference sampling campaigns conducted with taxonomic experts. The inclusion of a local reference library enabled us to significantly enhance metabarcoding efficiency. This improvement occurred in one of the world’s most thoroughly researched areas for beach meiofauna, and we expect that the impact in other regions and habitats will be even greater.

### Improved metabarcoding efficiency through enhanced molecular reference databases

We show that with a local reference library, 11,361,563 instead of just 4,633,286 sequences could be annotated to meiofauna, an increase of 6,728,277 sequences (145%) that would otherwise remain unassigned. On the genus and species level, reference barcoding increased the number of identified OTUs by over 200%. We identified more OTUs at all tidal levels, from dunes to the low tide line, and showed that patterns of community dissimilarity between tidal levels become more clear after adding new reference barcodes. Spatial analysis revealed that the enhanced database improved OTU assignment across all tidal levels. Our results of a clear separation in community composition between supralittoral areas of the beach, and areas in the lower intertidal zone, with samples from the swash zone around the high tide line falling in between, is in line with previous findings on beach meiofauna (Pereira et al., 2017). Our results show, however, that this pattern was more evident after adding new reference barcodes. We show that enhancing molecular reference databases significantly boosts metabarcoding efficiency for meiofauna across various ecological niches and geographic regions. While the advantages of enriched reference databases are documented for diverse ecosystems and taxonomic groups (Garlasché et al., 2023; Gold et al., 2021; Kjærandsen, 2022; Kocher et al., 2017; Magoga et al., 2022)), our results show a substantial improvement even in one of the most extensively studied areas for beach meiofauna. This underscores the massive benefit of local reference database enhancement on metabarcoding studies, underlining its value regardless of the prior level of research intensity.

### Combining morphological and molecular techniques

Our study emphasises the need for continuous efforts in combining traditional taxonomic identification and DNA barcoding, especially targeting under-represented regions and taxa, as the lack of taxonomic expertise and sampling leaves many beach meiofauna species undescribed (Martinez et al., 2023)(Curini-Galletti et al., 2012, 2023; Martínez et al., 2019). This hampers more detailed ecological studies that would facilitate the understanding of beach biodiversity and ecosystem processes, a task urgently needed due to increasing pressure on the globally important ecosystems (Lansu et al., 2024; Schlacher et al., 2007).

Combining morphological identification with molecular methods is increasingly feasible due to the development of rapid DNA barcoding pipelines relying on Oxford Nanopore technology (Srivathsan et al., 2019, 2023), potentially allowing the production of molecular references in field locations (Chang et al., 2020; Marin et al., 2022), while taxonomic experts are sorting and identifying specimens. An increased identification efficiency for metabarcoding data will significantly contribute to more precise ecological results (Faria et al., 2018), which will facilitate the use of molecular methods in monitoring (Aylagas et al., 2016), but also species detection (Giribet et al., 2023), and can therefore in turn facilitate taxonomic work on meiofauna.

### Change in inferred ecological patterns

We found that including new reference barcodes resulted in an increased identified OTU richness across all tidal levels, with the most significant increase observed in the middle intertidal zone. This led to a pattern of OTU richness peaking in the middle intertidal zone, which was more pronounced when annotating metabarcoding data with the NCBI reference database. This pattern, characterised by the highest richness in the middle intertidal zone and lower richness toward both the upper intertidal and supralittoral zones as well as the lower intertidal zone, thereby forming a bell-shaped diversity curve, has been previously described in studies based on morphological analyses of meiofauna (Armonies & Reise, 2000; Gingold et al., 2010; Maria et al., 2013). Thus, an improved reference database might reveal meiofauna richness patterns that align more closely with those identified through thorough, but time-intensive and expertise-dependent studies on morphology.

We found that the inclusion of new reference barcodes for annotating metabarcoding data increased the community dissimilarity between tidal levels, and led to a more distinct inferred community composition of the intertidal communities as opposed to communities from the supralittoral and the swash zone. This is likely because the addition of new reference barcodes allows for both the identification of more widespread meiofauna OTUs, which could not be identified with less complete reference databases, and rare OTUs found in only few sites. For example, OTU 1 in our dataset, present in 119 samples and with > 2 million reads, was only identified as a meiofaunal copepod after adding the new reference barcodes, and was unassigned based on annotation with NCBI. The meiofauna community composition in the middle to lower intertidal zones of the studied beaches may be more similar, potentially due to similar ecological conditions across these areas. In contrast, the higher variability between communities from the supralittoral levels and the swash zone, may indicate these areas’ exposure to more fluctuating environmental conditions or disturbances, leading to reduced community stability. This is plausible as the supralittoral zone of beaches is often heavily influenced by human activities, such as tourism-related activities including trampling and beach driving, which has been shown to have major impacts on meiofauna communities (Gheskiere et al., 2005; Martínez et al., 2020; Pereira et al., 2017)

### Limitations and biases in reference barcoding

Despite the major improvements in databases and metabarcoding efficiency, our study has limitations. The focus on a limited geographic region (here: the Netherlands’ west coast) for reference barcoding can introduce bias in the molecular reference database, as shown by our analyses on the correlation of increase in annotation efficiency with distance to the nearest sampling site used for reference barcoding. However, existing reference databases are almost always biassed towards well-studied taxonomic groups and regions, and are thereby inherently biassed (H. Weigand et al., 2019), a fact that is usually not tested due to the lack of comparison data from the same study. Our analyses show that the existing reference data in NCBI GenBank allows identifying the general patterns in meiofauna diversity in the study area, but the additional reference barcodes allow identifying more taxa and detect clearer patterns of alpha and beta diversity across the intertidal zone. In addition, the ‘local’ reference library provided significant improvements along 650 km of coastline and up to 300km away from the sites where the organisms were obtained for reference barcoding. Furthermore, primer bias can hamper the amplification of both new reference barcodes and taxa in metabarcoding datasets. This is a known phenomenon that affects several taxonomic groups that are numerous in beach meiofauna communities, such as Nematoda (Ren et al., 2023) and Platyhelminthes (Balsamo et al., 2020). However, the highly degenerate LerayXT primers allow for amplification of a wide range of meiofaunal taxa (J.-N. Macher et al., 2022), and we show that targeted reference barcode sequencing of individual meiofaunal specimens on Oxford Nanopore platforms can overcome this impediment, as we successfully amplified reference barcodes for all major meiofauna taxa.

### Future collaborative efforts for reference database expansion

Future research efforts should target both unexplored areas and taxa to enhance the taxonomic and geographical scope of reference databases, but also revisit areas that have been studied, as we unexpectedly found that COI reference barcodes for beach meiofauna in the North Sea region are rare. This gap is surprising because the North Sea has been the focus of extensive meiofauna studies (Armonies, 2018; Reise & Ax, 1979; Vincx et al., 1990). However, in general, only a limited number of studies on meiofauna incorporate molecular techniques, and there is a tendency to use the 18S rRNA marker due to its higher success rate in amplification, even though it offers less specificity in species identification (Gielings et al., 2021).

Our work demonstrates the feasibility of using COI barcoding and metabarcoding for a diverse range of beach meiofauna. The mitochondrial COI gene is the standard molecular barcode for most animals, and often offers superior species-level identification compared to 18S rRNA (Fontaneto et al., 2015; Giebner et al., 2020; Tang et al., 2012). Improving the 18S rRNA reference library is important, but might not be as helpful in determining meiofaunal species due to the conserved nature and thereby lower variability in this marker. It is therefore desirable to increase the availability of COI barcodes for meiofauna, as this will facilitate species-level molecular studies and enable more direct comparisons with macrofauna and other invertebrate groups, thanks to the use of the same amplified marker gene and gene region.

We advocate for the replication of our approach of combining taxonomic expert workshops, conducting reference barcoding, and applying metabarcoding to study meiofauna, but also other understudied taxonomic groups. Future efforts should focus on increasing the number of reference specimens that are identified at the species level. Additionally, integrating environmental data with metabarcoding results will offer insights into the ecological factors influencing meiofauna distribution and diversity.

## Conclusion

Our study highlights the benefits of enhancing molecular reference databases for DNA metabarcoding analyses through integrated taxonomy and DNA barcoding. This approach significantly improves species identification and biodiversity assessment, and shows the need for a collaborative effort in merging traditional taxonomic with molecular approaches. We advocate for continued efforts to build comprehensive, globally representative reference databases for meiofauna, which will be fundamental to advancing our understanding of coastal biodiversity and ecosystem dynamics.

## Acknowledgements

We would like to thank Elza Duijm, Nafiesa Ibrahim and Maarten van der Salm for support and help in the laboratory, Hannco Bakker and Tim Rietbergen for support with collection management, Kevin Beentjes for facilitating the sequencing, and Giada Spagliardi for help during the taxonomic workshops.

## Data Accessibility and Benefit-Sharing

All raw data is available in the NCBI Sequence Read Archive (SRA), under BioProject number PRJNA1081920. The reference specimen list is available as a GBIF dataset: https://doi.org/10.15468/gemfv4.

## Author Contributions

JNM, AM, WR, DF developed the study. All authors participated in sampling and identification of meiofauna during the taxonomic expert workshops and identified specimens. JM took samples for metabarcoding. JM, LVG DDJ performed metabarcoding lab work. JNM, AM, PEC, DF led the data analyses. JNM and DF led the writing. All authors approved the final version of the manuscript.

